# Pharmacologic Inhibition of DYRK1A Results in MYC Hyperactivation and ERK Hyperphosphorylation rendering *KMT2A*-R ALL Cells Sensitive to BCL2 Inhibition

**DOI:** 10.1101/2022.10.02.510349

**Authors:** Christian Hurtz, V. S. S. Abhinav Ayyadevara, Gerald Wertheim, John A Chukinas, Joseph P Loftus, Sung June Lee, Anil Kumar, Rahul S Bhansali, Srividya Swaminathan, Huimin Geng, Thomas Milne, Xianxin Hua, Kathrin M Bernt, Thierry Besson, Junwei Shi, John D. Crispino, Martin Carroll, Sarah K Tasian

## Abstract

*KMT2A*-rearranged (*KMT2A*-R) B cell acute lymphoblastic leukemia (ALL) is a high-risk disease in children and adults that is often chemotherapy resistant. To identify non-cytotoxic approaches to therapy, we performed a domain-specific kinome-wide CRISPR screen in *KMT2A*-R cell lines and patient derived xenograft samples (PDX) and identified dual-specificity tyrosine phosphorylation-regulated kinase 1A (DYRK1A) as a potential target. Pharmacologic inhibition of the KMT2A-fusion transcriptional co-regulator Menin released the KMT2A-fusion complex from the DYRK1A promoter thereby lowering DYRK1A expression levels confirming DYRK1A as a direct target of the KMT2A fusion oncogene. Direct pharmacologic inhibition of DYRK1A decreased cell proliferation of *KMT2A*-R ALL, thereby confirming the requirement of DYRK1A in this ALL subtype. To further understand the biologic function of DYRK1A in *KMT2A*-R ALL, we leveraged pharmacologic DYRK1A inhibitors in *KMT2A*-R PDX and cell line models. DYRK1A inhibition consistently led to upregulation of MYC protein levels, and hyperphosphorylation of ERK, which we confirmed via *in vivo* treatment experiments. Furthermore, DYRK1A inhibition decreased ALL burden in mice. Our results further demonstrate that DYRK1A inhibition induces the proapoptotic factor BIM, but ERK hyperphosphorylation is the driving event that induces cell cycle arrest. In contrast, combined treatment of *KMT2A*-R ALL cells *in vitro* and *in vivo* with DYRK1A inhibitors and the BCL2 inhibitor, venetoclax, synergistically decreases cell survival and reduced the leukemic burden in mice. Taken together these results demonstrate a unique function of DYRK1A specially in *KMT2A*-R ALL. Synergistic inhibition of DRYK1A and BCL2 may provide a low-toxic approach to treat this high risk ALL subtype.

## Introduction

*KMT2A*-rearranged (*KMT2A*-R) or mixed lineage leukemia rearranged (*MLL*-R) acute lymphoblastic leukemia (ALL) is the most common ALL subtype in infant ALL with a frequency of 70% and is also common in children and adults with a frequency of about 10%^1–3^. *KMT2A*-R ALL is a high-risk disease with a poor clinical outcome with a survival rate of <75% in children and <35% in infants and adults. Consolidation approaches, such as allogeneic stem cell transplant and chimeric antigen receptor T cell (CART cell) therapy, used in older children and adults are frequently not feasible in infants^4^. Therefore, we focused on identifying new targetable vulnerabilities in this high-risk ALL subset.

The use of small molecule inhibitors targeting activated and disease driving kinases has demonstrated significant clinical success. For instance, targeting the *BCR-ABL1* oncogene in patients with Philadelphia chromosome positive (Ph+) ALL has significantly improved the overall survival of patients^5,6^. Furthermore, BTK inhibition with ibrutinib significantly improved the survival of patients with in chronic lymphocytic leukemia compared to ofatumumab^7^ alone and FLT3 inhibition has been demonstrated to improve the survival of patients with FLT3 mutated acute myeloid leukemia (AML)^8^. Moreover, non-kinase targets are also under clinical investigation; small molecule inhibitors have been developed to target the interaction of the *KMT2A* fusion with Menin or the enzymatic activity of DOT1L, two molecules that are required for the transcriptional and epigenetic regulation of *KMT2A*-R target genes. However, while *in vitro* studies demonstrated promising results, initial *in vivo* studies are either not yet reported (Menin)^9–12^, or have not validated a consistent and sustained anti-leukemia benefit (DOT1L)^13^. Thus, continued improvement in therapeutic approaches is needed.

Given the success for small molecule inhibitors in some leukemias and the overall increased specificity of kinase inhibitors, we performed a kinome-wide domain specific CRISPR screen in 3 high-risk subtypes of ALL and found that dual-specificity tyrosine phosphorylation-regulated kinase 1A (DYRK1A) is specifically required for the survival of KMT2A-R ALL. DYRK1A is a dual specificity kinase that autophosphorylates a tyrosine residue for activation^14^ but otherwise primarily targets serine/threonine residues on substrates for phosphorylation. The protein has a proposed but incompletely defined role in cell cycle regulation that may vary depending on cell type. Significant research has characterized the role of DYRK1A in neuronal development and a smaller body of work has suggested its role in leukemia^15^. The DYRK1A gene is located on chromosome 21 which is triplicated in Down syndrome (DS, trisomy21). DS patients are at higher risk of developing acute megakaryoblastic leukemia and acute lymphoblastic leukemia, the pathophysiology of which may in part be related to *DYRK1A* overexpression^16,17^

Herein, we demonstrate an unanticipated direct regulation of DYRK1A via oncogenic *KMT2A*-rearrangements and a specific requirement of DYRK1A for normal proliferation of this high-risk ALL subtype. We also found that DYRK1A is required to suppress the pro-apoptotic molecule BIM and that DYRK1A inhibition consequently renders *KMT2A*-R ALL cells sensitive to BCL2 inhibition. Overall, these data provide new insights into the mechanisms of leukemogenesis by *KMT2A* fusion proteins and suggests a novel therapeutic approach for these leukemias.

## Materials and Methods

### Primary human ALL specimens and cell lines

Diagnostic bone marrow specimens from children and adults with ALL were obtained via informed consent on Institutional Review Board-approved research protocols of the Children’s Hospital of Philadelphia (CHOP) and University of Pennsylvania (UPenn) in accordance with the Declaration of Helsinki as previously described ^18,19^. Human KMT2A-R and non-KMT2A-R ALL cell lines were obtained via collaborators or purchased from the DSMZ biorepository (Braunschweig, Germany), validated by short tandem repeat analysis, and confirmed as *Mycoplasma*-free every 6 months^20–23^.

Human ALL cells harvested from murine spleens of some PDX models were cultured on OP9 stroma in Minimum Essential Medium (MEMα; Life Technologies) with GlutaMAX containing 20% FBS, 100 IU/mL penicillin, 100 μg/mL streptomycin, and 1 mM sodium pyruvate. Cell lines were cultured in Roswell Park Memorial Institute medium (RPMI; Life Technologies) with GlutaMAX containing 10 or 20% FBS, 100 IU/mL penicillin, 100 μg/mL streptomycin (hereafter referred to as ‘B cell medium’) at 37°C in a humidified incubator with 5% CO2 and maintained in culture for fewer than 3 months to minimize infectious contamination and genetic drift.

### Patient-derived xenograft modeling and *in vivo* preclinical drug trials

All PDX model establishment and experimental animal studies were conducted on Institutional Animal Care and Use Committee (IACUC)-approved research protocols at CHOP and UPenn in accordance with NIH and American Veterinary Association Guidelines for the Euthanasia of Animals. Briefly, KMT2A-R and non-KMT2A-R ALL PDX were created via intravenous injection of primary human ALL cells into 6–8-week-old male and female NOD/SCID/IL-2rγ^null^ (NSG), mice as previously described^18,19,24,25^. *In vivo* studies including human luciferase transduced cells were performed as previously described^26^. After flow cytometric confirmation of >5% CD45+ CD19+ PDX ALL in peripheral blood or via bioluminescent, animals were randomized to treatment with indicated combinations of (1) oral (PO) or intraperitoneal (IP) gavage vehicle and control chow, (2) harmine 10mg/kg or 20mg/kg (5x/week) or 20mg/kg rodent chow continuously provided, (3) venetoclax 50mg/kg (5x/week) or 10mg/kg PO once daily (3x/week), (4) for up to 10-20 days depending on rate of leukemia progression in control animals requiring study termination at a pre-determined endpoint^18,19,27^. Harmine hydrochloride-enriched chow was ordered via Research Diets, Inc.

### Flow cytometry analysis

Human and mouse leukemia samples were stained with cell surface antibodies according to manufacturer’s instructions using Fc block and respective isotype controls. Annexin V, propidium iodide, 7-AAD, or DAPI (BD Biosciences) were used for apoptosis analyses. EDUFlow cytometry was performed using FACSVerse or LSRII flow cytometers (BD Biosciences) with data analysis in Cytobank or FlowJo (TreeStar) and graphical display and statistical analysis in Prism (GraphPad); see Statistics below.

### Western Blotting

All cells were lysed in CelLytic buffer (Sigma) supplemented with 1% HALT (protease inhibitor and phosphatase inhibitor cocktail; Thermo Scientific). Protein samples were loaded on NuPAGE 4-12% Bis-Tris gradient gels (Invitrogen) and transferred on PVDF membranes (Millipore). For detection of mouse and human proteins, primary antibodies were used in combination with anti-rabbit or anti-mouse HRP-linked secondary antibodies (Cell Signaling) and Amersham ECL Western blotting detection reagent (GE Life Sciences).

### Cell proliferation and viability assays

One hundred thousand human ALL cells were seeded in a volume of 100 μL B cell medium/well (as described above) on Optilux 96-well plate (BD Biosciences, San Jose, CA). EHT1610, harmine, harmine hydrochloride, and venetoclax were diluted in medium and added at the indicated concentration in a total culture volume of 150 μL. MI-503 was synthesized by Wuxi Pharma as previously described^11,12,63^. After culturing for 72 hours, cell proliferation and viability were measured by XTT (Cell Signaling) according to the manufacturer’s instructions with fluorescence read at 450 nm. Fold changes were calculated using baseline values of untreated cells as a reference (set to 100%) and displayed graphically in Prism.

### Quantitative real-time polymerase chain reaction (qRT-PCR)

Total RNA from leukemia cells was extracted using RNeasy isolation kit (Qiagen). cDNA was generated using the qScript cDNA SuperMix (Quanta). Quantitative real-time PCR was performed with Power SYBR^®^ Green Master Mix (Applied Biosystems) and 7500Real Time PCR System (Applied Biosystems) according to standard PCR conditions.

### In vitro drug synergy analysis

The expected drug combination responses between EHT1610 and trametinib or EHT1610 and venetoclax were calculated based on the ZIP reference model using SynergyFinder^64^. Deviations between observed and expected responses with positive and negative values denote synergy and antagonism, respectively. For estimation of outlier measurements cNMF algorithm^65^ implemented in SynergyFinder was utilized.

### Retroviral and lentiviral transduction

Transfections of 293FT cells with retroviral and lentiviral constructs were performed using lipofectamine 2000 with Opti-MEM media (Invitrogen). Viral supernatants were produced by co-transfecting 293FT cells with the viral gag-pol and packaging vectors together with the Cas9 or sgRNA constructs. Cultivation was performed in high glucose Dulbecco’s modified Eagle’s medium (DMEM; Invitrogen) with GlutaMAX containing 10% fetal bovine serum, 100 IU/mL penicillin, 100 μg/mL streptomycin, 25mM HEPES, 1 mM sodium pyruvate, and 0.1 mM non-essential amino acids. Regular media were replaced after 16 hours by growth media containing 3 mM caffeine. After 24 hours, the viral supernatant was harvested and filtered through a 0.45 μm filter. Retroviral transductions were performed by loading the viral supernatant on 50 μg/mL retronectin-coated (Takara) non-tissue 6-well plates prior to centrifugation (2000 x g, 90 min at 32°C) two times. Subsequently, 1-2 × 10^6^ cells were loaded and transduced per well by centrifugation at 600 x g for 30 minutes and maintained for 72 hours at 37°C with 5% CO2 prior to transfer into culture flasks. Lentiviral transductions were performed by loading viral supernatant on 50 μg/mL retronectin-coated non-tissue 6-well plates with 1-2 x 10^6^ cells. Cells were centrifuged x 30 minutes at 600 x g and maintained for 16 hours at 37°C with 5% CO2 prior to replacing media with fresh RPMI and GlutaMAX containing 20% FBS, 100 IU/mL penicillin, and 100 μg/mL streptomycin. Cells were then cultured at 37°C with 5% CO2 prior to analysis.

### Pooled CRISPR Screening

The kinase domain-focused sgRNA library screening has been performed as previously described.^66^. To perform the kinome-wide kinase sgRNA screen, we first virally transformed SEM, HAL-01, and TVA one cells with a LentiV_Cas9_puro vector^66^. After successful puromycin selection, we then further transduced the ALL cells with lentivirus of pooled sgRNA library as described. Virus titer was measured by infection of cells with serially diluted virus. For transduction of single sgRNA per cell, multiplicity of infection (MOI) was set to 0.3-0.4. To maintain the representation of sgRNAs during screen, the number of cells was kept 1000 times more than sgRNA number in the library. Cells were harvested at initial (day 3 post-infection) and final (21 days after initial) timepoints. Genomic DNA was extracted using QIAamp DNA mini kit (QIAGEN). Analysis was performed as previously described^67^.

The secondary reduced library screen was performed on SEM cells as described above. The reduced library included 14 kinases that we have identified in our primary screen to be required for *KMT2A*-R ALL survival and 21 negative controls.

### Statistical analyses

Data display and statistical analyses were performed using Prism 7 (GraphPad). Significance values for *in vitro* and *in vivo* studies were calculated using unpaired two-tailed student t-test (2 groups) or ANOVA with Dunnett’s or Tukey’s post-test for multiple comparisons (3 or more groups) to determine statistical significance. Survival analysis distribution between 2 groups for clinical patient samples was determined by the log-rank test. A *P* value of less than 0.05 was considered significant. All values are expressed as mean ± SEM.

## Results

### KMT2A-R ALL is dependent on DYRK1A expression for their survival

We performed a domain-specific kinome-wide CRISPR screen in high-risk acute lymphoblastic leukemia cell lines (SEM and HAL-01) as well as a PDX sample (TVA-1; *ETV6::ABL1*^20^) to identify novel druggable targets in high-risk ALL **(Figure 1A)**. As expected, we identified dependencies upon 1) FLT3 in the FLT3 wild-type overexpressing SEM (*KMT2A::AFF1*) cell line, 2) the checkpoint inhibitors ATR and CHK1 in the HAL-01(*TCF3-HLF*) ALL cell line, and 3) ABL1 in immortalized TVA-1 cells, validating the specificity of our screens **(Figure 1B and Supplemental Figure 1A)**. SgRNAs targeting DYRK1A were specifically reduced in SEM cells compared to the other tested samples **(Figure 1C)**. Furthermore, DYRK1A is not a common essential gene validated via the Cancer Dependency MAP website indicating that targeting DYRK1A may represent a target with a unique therapeutic window **(Supplement Figure 1B)**. 5 Different DYRK1 family members are known and only DYRK1A was required for *KMT2A-AFF1* cell survival **(Figure 1D)**. The importance of DYRK1A was validated in a second CRISPR screen consisting of 14 kinases that we identified in our initial screen **(Figure 1E)**. Taken together, our results demonstrate a unique dependency of *KMT2A-AFF1* cells on DYRK1A in our screen.

**Figure 1:**
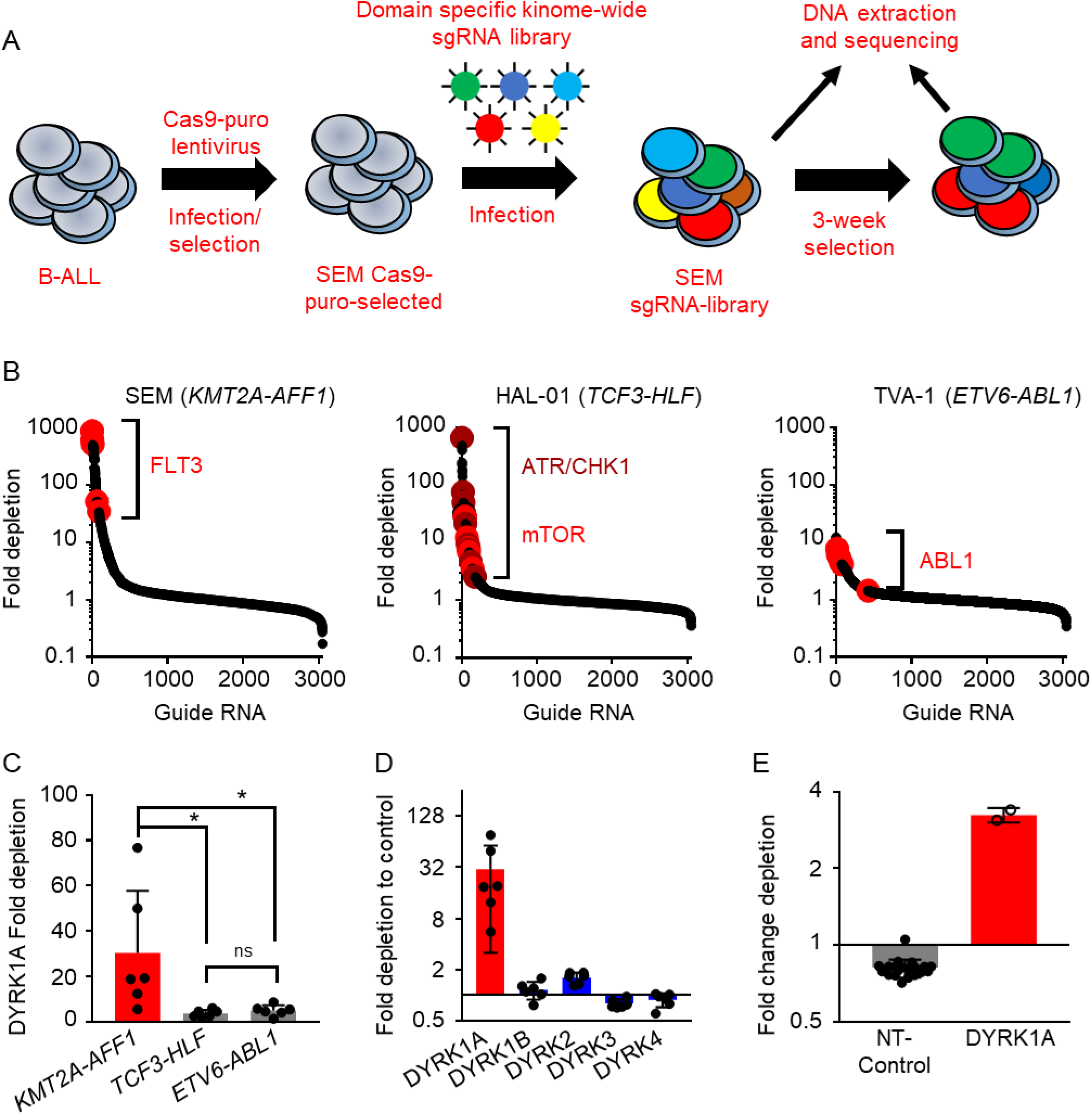
Domain-specific kinome-wide CRISPR screen in high-risk ALL. **A)** Overview of the performed domain specific kinome-wide library screen. SEM (*KMT2A::AFF1*), HAL-01 (*TCF3::HLF*), and TVA-1 (*ETV6::ABL1*) cells were virally transduced with a lentiviral Cas9-Puro construct. After puromycin selection, the cells were transduced with a GFP tagged domain specific kinome-wide CRISPR-library consistent of 550 kinases, 100 negative controls and 50 positive controls (6 guide RNAs (sgRNA) per kinase). Cells were collected at 2 time points for sequencing; 3 days after the library transduction, which served as input control and 3-weeks after the transduction which served as the experimental readout. **B)** Shown is the fold depletion of the individual sgRNAs in the tested samples. Highlighted are known positive controls for each individual ALL subtype. **C)** Shown is the fold depletion of DYRK1A among the different ALL subtypes. **D)** Shown is the fold depletion of each DYRK1A family member in SEM cells. **E)** Cas9-puro-selected SEM cells were lentiviral transduced with a smaller sgRNA-GFP library. The library included two sgRNAs targeting one of 14 previously identified targets including 21 negative controls.

### Oncogenic KMT2A fusion proteins directly bind to the DYRK1A promoter and regulate its expression

Given that our CRISPR-screen identified DYRK1A to be specifically important for *KMT2A*-R ALL, we hypothesized that oncogenic KMT2A fusions may directly regulate DYRK1A expression. Strikingly, analysis of previously published ChIP-Seq data of *KMT2A::AFF1* ALL cells demonstrate that both *KMT2A* and *AFF1* directly bind to the promoter region of DYRK1A in ALL cell lines SEM and RS4;11 **(Figure 2A)**^28,29^. For the ChIP-Seq experiment antibodies binding to the N-terminal region of *KMT2A* and the C-terminal region of *AFF1* were used. Given that *KMT2A*-R ALL is often a monoallelic balanced translocation with wild type *KMT2A* and *AFF1* still present in the cells, we then analyzed a ChIP-Seq data set that used human leukemia cells that were generated by overexpression of a *KMT2A-Aff1*-Flag construct. Using a Flag antibody allowed for identification of oncogene specific binding partners. Strikingly, *KMT2A-Aff1* directly bound to the DYRK1A promoter region, validating that KMT2A fusions directly binds to the DYRK1A promoter region **(Figure 2B)**. The binding of the KMT2A fusion construct to MYC is shown as control.

**Figure 2:**
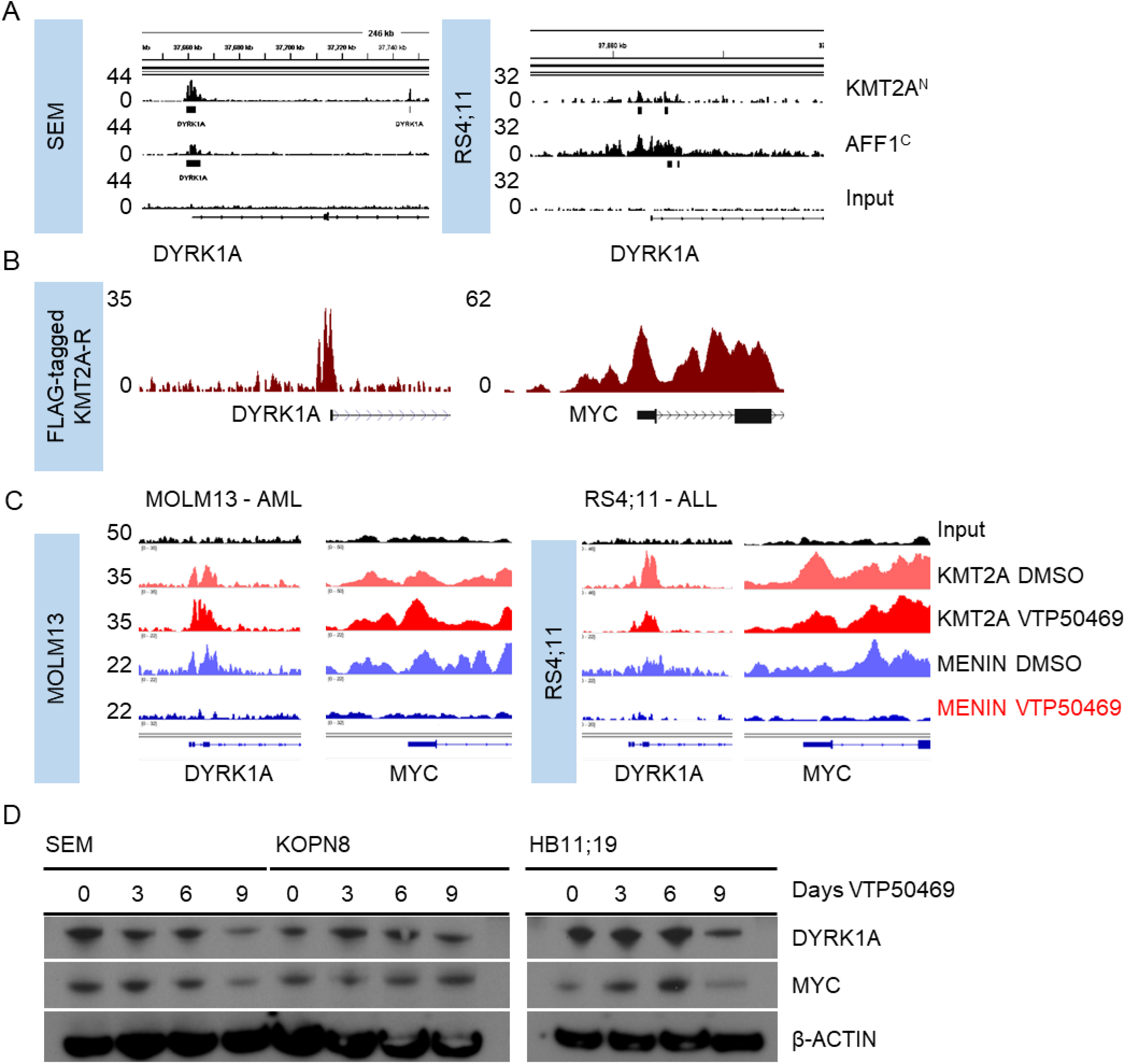
Oncogenic *KMT2A*-rearrangements transcriptionally regulate DYRK1A. **A)** ChIP-Seq (Chromatin immunoprecipitation [ChIP] combined with high-throughput sequencing) tracks on the DYRK1A promoter region using an antibody specific for the *KMT2A* N- and AFF1 C-terminal domain were used. The Y-axis represents the number of reads for peak summit normalized by the total number of reads per track. Data from two *KMT2A-AFF1* cell lines SEM (GSE83671)^28^ and RS4;11 (GSE38403)^29^ are shown. **B)** ChIP-Seq tracks on the DYRK1A promoter region using a FLAG-specific antibody in *KMT2A*-*Aff1*-FLAG-transformed human ALL cells. The Y-axis represents the number of reads for peak summit normalized by the total number of reads per track (GSE84116)^56^. **C)** ChIP-Seq tracks on the DYRK1A and MYC promoter region using *KMT2A* N-terminal and Menin specific antibodies. Two *KMT2A*-R cell lines (MOLM13/AML and RS4;11/ALL) were either treated with control or the Menin inhibitor VTP50469 as previously described (GSE127508)^30^. **D)** Western blot analysis of DYRK1A, MYC and β-actin in three *KMT2A*-R ALL cells (SEM, KOPN8, RS4;11) treated with the menin inhibitor VTP50469 (100nM) for the indicated days.

To further validate if KMT2A fusions not only bind to the promoter region, but also directly affect DYRK1A transcription, we next analyzed ChIP-Seq data from the *KMT2A::MLLT3* acute myeloid leukemia (AML) cell line MOLM13 and RS4;11^30^. Cells were treated with vehicle or the Menin inhibitor VTP50469^30^ for 3 days. Strikingly, Menin inhibition prevented the binding of Menin, an important regulatory unit that is required for the transcriptional regulation of KMT2A fusions target genes, to the *DYRK1A* promoter. *MYC* was analyzed as positive control demonstrating reduced binding of Menin to the MYC promoter site **(Figure 2C)**. We next analyzed gene expression data of SEM cells treated either with the DOT1L inhibitor EPZ004777, the Menin inhibitor Mi-2-2 or a combination of both drugs. Strikingly, the analysis demonstrates that DYRK1A was specifically downregulated after Menin inhibition, while DOT1L inhibition only marginally reduced the DYRK1A expression levels **(Supplement Figure 1C)**. We conclude that KMT2A fusions transcriptionally regulate DYRK1A via menin, but not epigenetically via DOT1L.

To validate this finding, we treated the SEM cells with the Menin inhibitor, MI-503^12^ for 7 days. RT-PCR analysis demonstrated that Menin inhibition results in a significant reduction of DYRK1A as well as other known transcriptional targets **(Supplement Figure 1D)**. To validate that the regulation of DYRK1A is specific to the KMT2A fusion, not the wild type allele, we analyzed gene expression data of conditional *KMT2A* knock out mice after 3 days of *KMT2A* deletion and strikingly, genetic deletion of normal *KMT2A* did not affect the DYRK1A RNA levels **(Supplement Figure 1E)**. Taken together, these data demonstrate that DYRK1A transcriptional regulation is unique to the oncogenic KMT2A fusion.

### DYRK1A is highly expressed in KMT2A-R ALL and high expression levels are retained during relapse

We next tested the DYRK1A protein levels across different subtypes of ALL and AML and, interestingly, DYRK1A expression was increased in *KMT2A*-R ALL compared to the other tested cell lines **(Supplement Figure 1 F)**. Furthermore, we tested multiple PDX cases from *KMT2A*-R ALL patients at diagnosis (black) and relapse (red) and compared them to non-*KMT2A*-R ALL. The data demonstrates that the high DYRK1A protein levels are retained at relapse and are elevated in *KMT2A*-R ALL compared to Ph-like ALL **(Supplement Figure 1G)**. Finally, we performed a western blot analysis of different *KMT2A*-R ALL cell lines that were treated with the Menin inhibitor VTP50469 for 3, 6, and 9 days. MYC, a known target of *KMT2A*-R which is involved in cell proliferation was used as a positive control. Similar to what we have seen at the transcriptional level, the protein expression of DYRK1A was reduced after 9 days of Menin inhibition **(Figure 2D)**. However, menin inhibition did not fully abrogate the DYRK1A protein expression **(Figure 2D)** and only affected cell proliferation of *KMT2A*-R ALL cells after multiple days of menin inhibition (**Supplement Figure 2 A)**. Note that in these studies, apoptosis was only induced when high concentrations of VTP50469 were used **(Supplement Figure 2B)** consistent with previous results^30^. Overall, these data support that KMT2A fusions regulates DYRK1A RNA and protein expression levels via direct transcriptional regulation that requires menin binding.

### DYRK1A negatively regulates MYC

As observed above, KMT2A fusions directly regulate both *MYC* and *DYRK1A* expression. However, it has also been shown that viral overexpression of DYRK1A mediates MYC phosphorylation (Ser62 and Thr58) and consequently MYC degradation in non-*KMT2A*-R AML, suggesting that DYRK1A may negatively regulate MYC^31^. To further test if there is a KMT2A fusion independent regulation of MYC via DYRK1A in *KMT2A*-R ALL, we first tested if genetic deletion of *Dyrk1a* results in higher *MYC* RNA levels in B cells using a conditional *Dyrk1a* KO mouse model^17^. Strikingly, we confirmed that genetic deletion results in potent upregulation of Myc in *Dyrk1a-* deleted cells, indicating that Dyrk1a negatively regulates Myc **(Figure 3A)**. To validate this finding in *KMT2A*-R ALL at the protein level, we treated *KMT2A*-R ALL cell lines with the DYRK1A inhibitor EHT1610 without modifying KMT2A fusion activity^32^. As hypothesized, we detected potent MYC upregulation upon DYRK1A inhibition in all tested cell lines **(Figure 3B).** Moreover, analysis of gene expression data from *KMT2A*-R ALL patient samples (COG P9906)^33^ demonstrates that DYRK1A and MYC expression levels are negatively correlated **(Figure 3C)**. Based on these findings we conclude that DYRK1A regulates MYC levels independent of the *KMT2A*-fusion oncogene. To further elucidate if there is a regulatory feedback mechanism between DYRK1A and MYC, we analyzed DYRK1A expression levels at different stages during B cell development including transgenic mouse model systems with overexpression of *Bcl6*, Myc/Bcl6 or deletion of *Lig4/tp53*, three distinct models of B cell lymphoma^34^. Notably, tumor cells isolated from the *Bcl6* transgenic and the *Myc/Bcl6* transgenic mice displayed reduced *Dyrk1a* mRNA expression levels **(Figure 3D and Supplement Figure 3A)** indicating that Dyrk1a may be negatively regulated either by Bcl6 or Myc.

**Figure 3:**
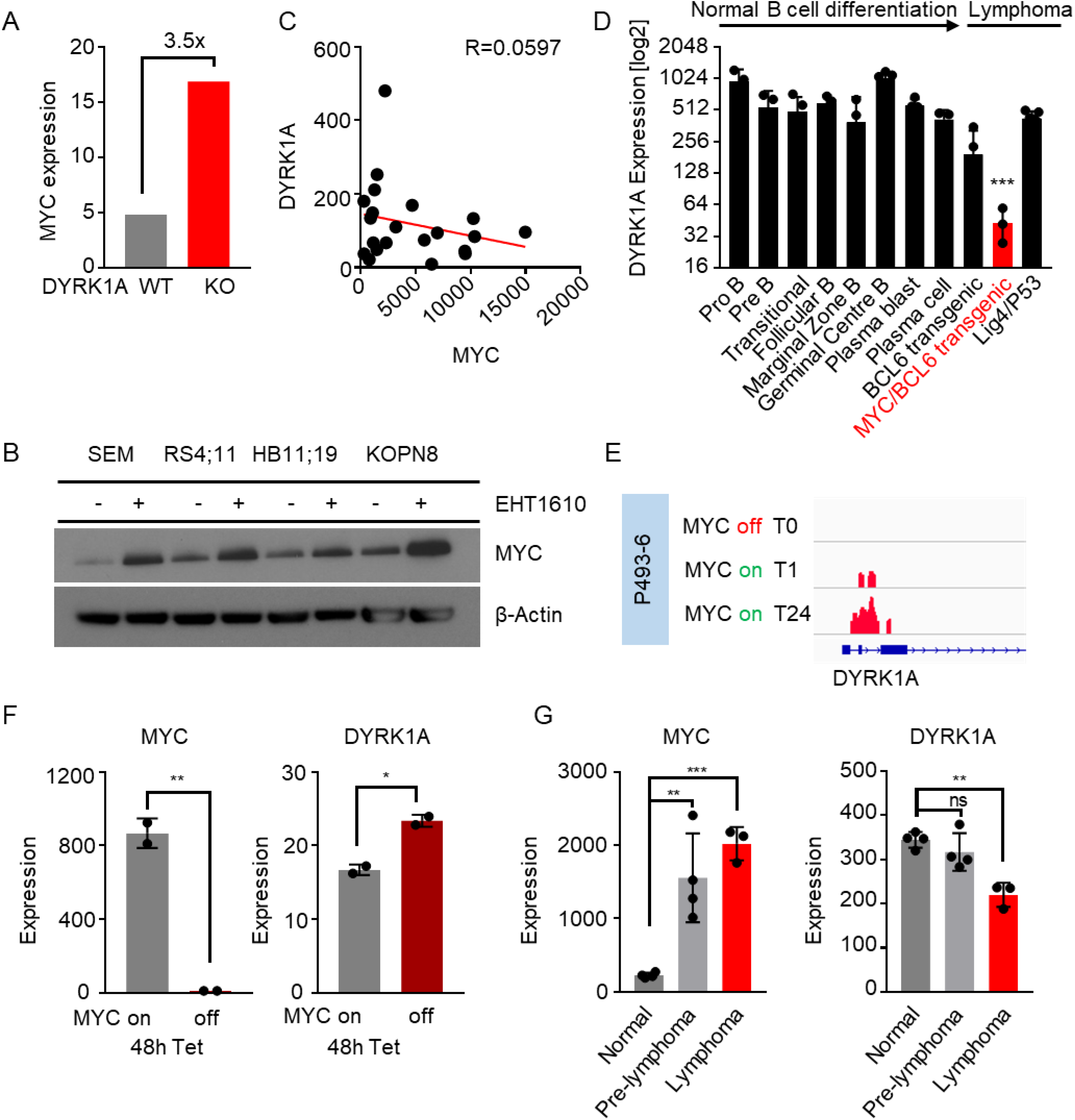
Feedback regulation between DYRK1A and MYC. **A)** MYC gene expression levels in DYRK1A wildtype and DYRK1A knock-out murine pre-B cells (GSE67052)^17^. **B)** Western blot analysis of MYC and β-actin in *KMT2A*-R ALL cell lines treated either with control or EHT1610 (5μM/72h). **C)** Correlation between the MYC and DYRK1A gene expression levels in *KMT2A*-R ALL pediatric patient samples from the Children’s Oncology Group (GSE11877)^33^. **D)** DYRK1A gene expression levels in comprehensive panel of purified developmentally defined normal murine B cells and genetically distinct murine lymphoma models (GSE26408)^34^. **E)** Chip-Seq experiment the MYC inducible Burkitt’s lymphoma cell line P493-6. Gene tracks of Myc binding at the DYRK1A promoter region at 0h (top), 1h (middle), and 24h (bottom)M are shown (GSE36354)^36^. **F)** MYC and DYRK1A gene expression levels in P493-6 cells before and after MYC inactivation. MYC inactivation was induced via tetracycline treatment for 48h (GSE120246). **G)** *MYC* and *DYRK1A* gene expression levels in Eμ-*myc* transgenic mice. These mice develop a fatal lymphoma within a few months of birth (GSE51008)^39,57^.

### MYC but not BCL6 negatively regulates DYRK1A

We next asked if BCL6 is a negative regulator of DYRK1A given its function as a transcriptional repressor^35^. We analyzed ChIP-Seq data from 2 ALL PDX cases (ICN12 and ICN13) as well as 3 cells lines (RCH-ACV, RS4;11, SEM) and identified that BCL6 does not directly bind to the *DYRK1A* promoter **(Supplement Figure 3B)**. Furthermore, genetic *Bcl6* deletion in BCR-ABL1 mouse ALL-like cells only resulted in moderate upregulation of Dyrk1a in one out of three probe sets **(Supplement Figure 3C)**. Induced BCL6 deletion in an ALL-like mouse knockout model system with overexpression of TCF3-PBX2 did not change the RNA expression levels of DYRK1A further validating that BCL6 does not regulate DYRK1A transcript levels **(Supplement Figure 3D)**.

To specifically test if MYC regulates DYRK1A transcriptionally, we analyzed ChIP-Seq data using the human P493-6 B cell model of Burkitt lymphoma^36^. Here MYC expression can be induced via removal of doxycycline form the cell culture medium^36^. Activation of MYC expression resulted in increased binding of MYC to the DYRK1A promoter in a time-dependent manner **(Figure 3E)**. To further test if MYC negatively regulates DYRK1A we performed a loss of function analyses by analyzing gene expression data of P493-6 before and after 48h of doxycycline-induced MYC deletion, which resulted in significant upregulation of DYRK1A **(Figure 3F)**. We validated this finding by analyzing BCR-ABL1 transformed ALL-like cells from a conditional *Myc* knock out mouse model system^37,38^ **(Supplement Figure 4)**. To further validate if Myc negatively regulates Dyrk1a, we performed a gain-of-function meta-analysis of a Eμ-myc transgenic lymphoma/leukemia model^39,40^ **(Figure 3G)**. The data demonstrate that the levels of MYC increase during the process of overt malignancy development. Strikingly, the levels of Dyrk1a are consistently reduced during the lymphoma development of these mice, further confirming a negative regulation of Dyrk1a via Myc.

Taken together, our analyses demonstrate that DYRK1A and MYC negatively regulate each other. *Dyrk1a* genetic deletion or pharmacologic inhibition results in increased MYC levels, while genetic deletion or overexpression of MYC mediate either the increase or decrease of DYRK1A, respectively.

### KMT2A-R ALL cells are sensitive to pharmacologic DYRK1A inhibition

To evaluate DYRK1A as a potential target for the treatment of ALL, we first tested if pharmacologic inhibition of DYRK1A induces apoptosis or cell cycle arrest selectively in *KMT2A*-R ALL *in vitro*. It has previously been demonstrated that DYRK1A is important for Down Syndrome associated ALL and the closely related *CRLF2-R* ALL^17,41^. We therefore excluded these ALL types from our analysis. We used 2 distinct DYRK1A inhibitors for our study 1) EHT1610 **(Figure 4A)** and 2) Harmine **(Supplement Figure 5A)**. Strikingly, DYRK1A inhibition with EHT1610, specifically reduced cell proliferation in *KMT2A*-R ALL compared to non-*KMT2A*-R ALL **(Figure 4B)**. We confirmed the sensitivity of *KMT2A*-R ALL using the DYRK1A inhibitors harmine and harmine hydrochlorate, **(Supplemental Figure 5B)**^42^. Based on the finding that DYRK1A significantly reduced the leukemia cell proliferation, we tested if pharmacologic inhibition of DYRK1A could be used as a therapeutic strategy. Preliminary work using EHT1610 in NSG mice showed toxicity and we therefore used harmine hydrochloride for these studies as an alternative DYRK1A inhibitor. We injected NSG mice with luciferase transduced SEM cells and treated the mice twice a day with 10mg/kg or 20mg/kg harmine hydrochloride for 10 days **(Figure 4C-D)**. Strikingly, bioimaging confirmed a potent leukemia reduction in the DYRK1A inhibitor treated group **(Figure 4C)**. Mice were sacrificed after the last treatment on day 10, splenocytes were purified and assessed for MYC induction **(Figure 4D)**. MYC protein expression was induced confirming that harmine at both concentrations inhibited DYRK1A. Taken together, these data demonstrate DYRK1A is required for *KMT2A*-R ALL cell proliferation.

**Figure 4:**
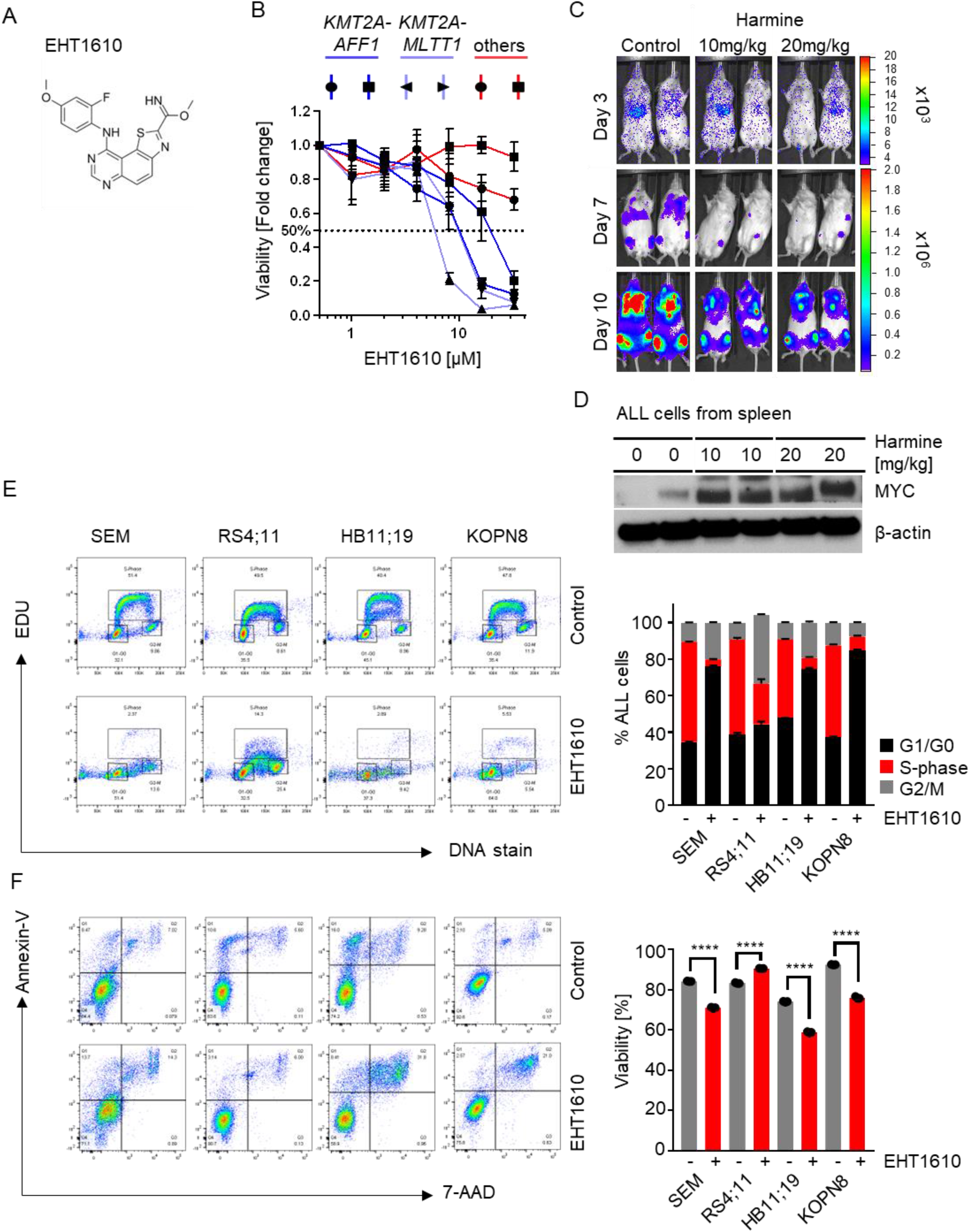
DYRK1A is required for *KMT2A*-R ALL cell proliferation. **A)** Molecular structure of EHT1610^32^. **B)** *KMT2A-AFF1* (dark blue), *KMT2A*-MLTT1 (light blue) and non-*KMT2A*-R ALL cell lines were treated with increasing concentrations of EHT1610. Viability/cell proliferation was determined via XTT assay after 72h. **C)** NSG mice were injected with firefly luciferase transduced HB11;19 cells. After 72h mice were treated twice a day either with 1) control or with harmine hydrochloride at the indicated concentrations. Bioimages were taken after 3, 7, and 10 days. **D)** HB11;19 cells from were isolated from mouse spleens 10 days after the indicated treatments in (C) and protein levels of MYC and β-actin were determined via Western blotting. **E)** Cell cycle analysis of the indicated *KMT2A*-R ALL cell lines after treatment with 5μM EHT1610 for 72h. On the left are representative examples of the flow cytometric analysis (n=3) and on the right is the summary of all three experiments. F) Cell viability and apoptosis was tested in control- and DYRK1A inhibitor-treated (EHT1610; 5μM; 72h) *KMT2A*-R ALL. On the left are representative examples of the flow cytometric analysis (n=3) and on the right is the summary of all three experiments.

### DYRK1A inhibition significantly reduced the S-phase of KMT2A-R ALL cells

It has been described in B- and T-cells that DYRK1A inhibition results in an accumulation of cells in S-Phase. To evaluate if DYRK1A inhibition affects cell proliferation or induces apoptosis, we first performed a cell cycle analysis before and after EHT1610 treatment, which demonstrated a significant reduction of cells in S-phase **(Figure 4E)**, while the cell viability was only marginally affected **(Figure 4F)**. Taken together, DYRK1A inhibition specifically affects cell proliferation by increasing the number of cells in G1/G0 and limiting the number of cells in S-phase in *KMT2A*-R ALL.

### DYRK1A represses ERK signaling and protects KMT2A-R ALL cells from apoptosis

It has been demonstrated that DYRK1A overexpression results in increased ERK signaling output in varied cell types^43–45^. Based on these findings we hypothesized that the induction of cell cycle arrest in *KMT2A*-R ALL cells after pharmacologic inhibition may be via DYRK1A inhibitor mediated down regulation of ERK signaling. To assess the effect of DYRK1A inhibition on ERK signaling, we treated different *KMT2A*-R ALL cell lines with the EHT1610 and collected protein samples at 4 different timepoints. Surprisingly, our results demonstrate that DYRK1A inhibition resulted in hyperphosphorylation of ERK in two out of the three samples tested **(Figure 5A)**. ERK hyperphosphorylation was confirmed using harmine and harmine hydrochloride in 3 out of 4 tested *KMT2A*-R ALL cell lines **(Supplement Figure 5C)**. To validate if this is a specific affect in *KMT2A*-R leukemias or if DYRK1A also mediates hyperphosphorylation in other ALL subtypes, we performed a Western blot to test the ERK phosphorylation levels in *KMT2A*-R ALL and non-*KMT2A*-R ALL cell lines as well as *KMT2A*-R acute myeloid leukemia cell lines either treated with control or EHT1610. Strikingly, ERK hyperphosphorylation was restricted to *KMT2A*-R ALL samples **(Figure 5B)**. It has been demonstrated that hyperactivation of B cell receptor signaling molecules can result in negative selection of B cells^46^. Based on these findings we hypothesize that ERK hyperphosphorylation may result in negative selection and cell cycle arrest in *KMT2A*-R ALL cells. To confirm this hypothesis, we first treated 4 different *KMT2A*-R ALL cell lines either with control, EHT1610, the MEK1/2 inhibitor trametinib, or a combination of both **(Figure 5C)**. As expected, DYRK1A inhibition resulted in potent ERK hyperphosphorylation in 3 out of the 4 tested *KMT2A*-R ALL cell lines, while trametinib inhibited ERK signaling. Most importantly, trametinib effectively prevented the EHT1610 mediated hyperphosphorylation of ERK. To test if trametinib would rescue *KMT2A*-R ALL cell proliferation, we performed a synergy study using increasing concentrations of EHT1610 and trametinib. Strikingly, our results demonstrate that trametinib rescued *KMT2A*-R ALL cells from DYRK1A inhibition induced cell death.

**Figure 5:**
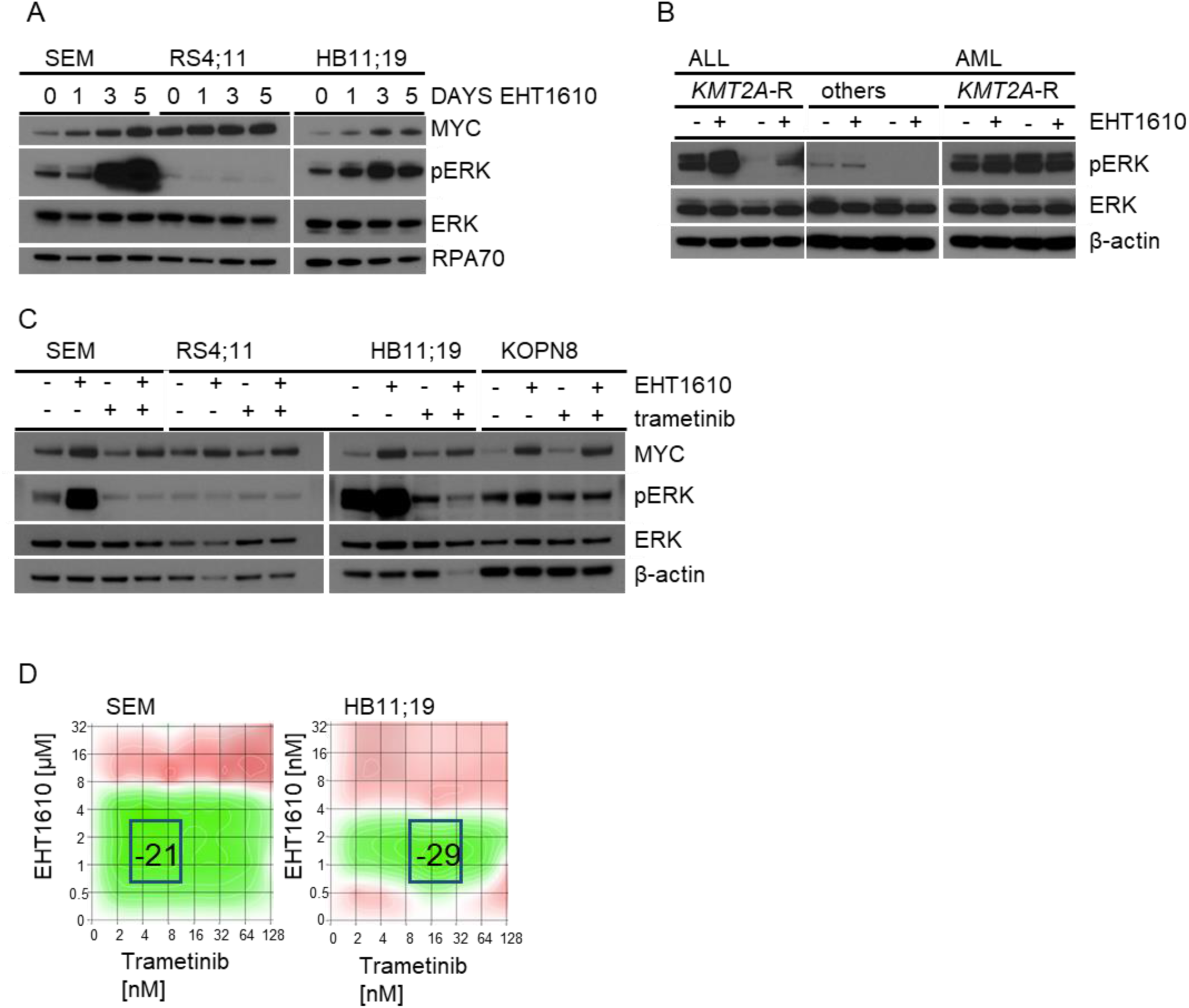
DYRK1A induces cell cycle arrest via ERK mediated signaling pathway hyperactivation. **A)** Protein expression levels of the indicated proteins were determined via Western blot using three *KMT2A*-R ALL cell lines treated with 5μM EHT1610 for the indicated time points. **B)** *KMT2A*-R and non-*KMT2A*-R ALL cell lines as well as *KMT2A*-R AML cell lines were treated either with control or 5μM EHT1610. After 72h pERK, ERK and β-ACTIN protein levels were determined via Western blot. **C)** The indicated *KMT2A*-R ALL cell lines were treated either with control, 5μM EHT1610, 20nM trametinib, or a combination of both. After 72h the expression levels of the indicated proteins were determined via Western blotting. **D)** The combinatorial effect of EHT1610 and trametinib was determined via synergy analysis.

Taken together, our results demonstrate an unexpected involvement of DYRK1A to regulate the ERK signaling output in *KMT2A*-R ALL cells and illustrates that DYRK1A inhibitor-mediated cell cycle arrest is induced via ERK hyperphosphorylation.

### DYRK1A inhibition renders cells sensitive to venetoclax

The proapoptotic factor BIM is regulated via multiple regulatory mechanisms. One of these regulators is MYC, which has been demonstrated to directly bind to the BIM promoter region to activate BIM mediated apoptosis^47^. The signaling molecule ERK has also been studied as a molecule that can either increase the proapoptotic activity via phosphorylation of Ser44, Thr56, and Ser58 (BIM Exon3)^48^ or decrease the BIM protein stability and induce proteasome-dependent degradation via phosphorylation of Ser69 (BIM Exon3)^49,50^. To evaluate if DYRK1A inhibition and consequent activation of MYC and ERK signaling results in increased BIM expression and activity in *KMT2A*-R ALL cells, we treated 4 cell lines with and performed a Western blot analysis of BIM and BCL2. The antiapoptotic activity of BIM is neutralized when BCL2 binds to BIM^51^. Strikingly, DYRK1A inhibition increased BIM expression in three of the four *KMT2A*-R ALL cell lines, while the BCL2 levels remained mostly unchanged **(Figure 6A)**. To determine if DYRK1A inhibition sensitizes *KMT2A*-R ALL cells to BCL2 inhibition, we treated *KMT2A*-R ALL cells with increasing concentrations of EHT1610 and the BCL2 inhibitor venetoclax. Strikingly, our data indicates that both drugs synergistically kill *KMT2A*-R ALL cells **(Figure 6B)**. To validate this finding, we treated *KMT2A*-R ALL cells with EHT1610, venetoclax, and a combination of both. Flow cytometric analysis confirmed that both drugs potently kill leukemia cells *in vitro* **(Figure 6C)**. Finally, we assessed the potential anti-leukemia activity of dual DYRK1A and BCL2 inhibition in a *KMT2A::AFF1* ALL PDX model. Indeed, DYRK1A inhibition resulted in potent reduction of the leukemia burden compared to control cells **(Figure 6D)**. However, BCL2 inhibition alone and in combination profoundly reduced the leukemia burden to undetectable levels. Only after 5 weeks of treatment, we detected a reoccurrence of the disease which was significantly lower in the dual treated group **(Figure 6E)**. Taken together, our results demonstrate that DYRK1A negatively regulates BIM. Furthermore, dual inhibition of DYRK1A and BCL2 synergistically killed *KMT2A*-R ALL cells in vitro and significantly reduced the leukemia burden *in vivo*.

**Figure 6:**
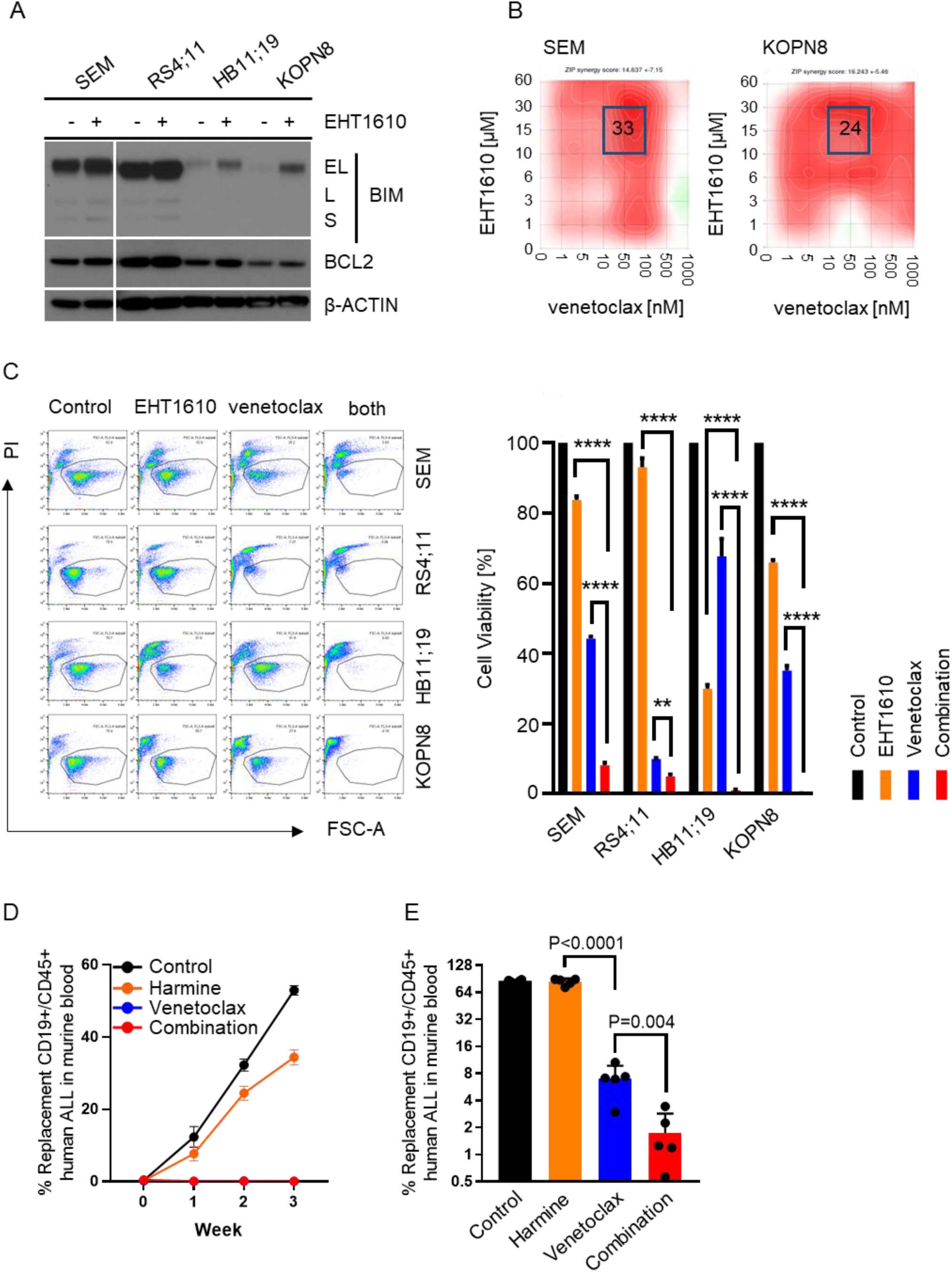
Dual DYRK1A and BCL2 inhibition synergistically kills *KMT2A*-R ALL cells. **A)** Western blot analysis of the indicated *KMT2A*-R ALL cell lines treated either with control or 5μM EHT1610. After 72h the expression levels of the indicated proteins was determined. **B)** SEM and KOPN8 cells were treated with increasing concentrations of EHT1610 and venetoclax. The synergistic effect of both drugs was determined via the Synergy Finder software as in Figure 5. **C)** 4 *KMT2A*-R ALL cells were treated with 5μM EHT1610 and 20nM venetoclax for 72h. Shown are representable examples of the flow cytometric viability analysis (left) and the statistical analysis right; n=3). **D)** *KMT2A*-R PDX mice (model ALL3113) were randomized to treatment with vehicle (control), 20mg/kg harmine via rodent chow, 50mg/kg venetoclax via PO injections once daily, or both inhibitors for the indicated time points (n=5). Human ALL progression was monitored via flow cytometric analysis of CD19+/CD45+ human ALL cells in murine blood over time.

## Discussion

To identify new targets in *KMT2A*-R ALL for potential precision medicine therapeutic approaches, we performed a kinome-wide CRISPR screen and identified multiple kinases required for the survival of *KMT2A*-R ALL cells. We selected to further study the importance of DYRK1A for *KMT2A*-R ALL as it met the following three criteria: 1) Growth inhibition upon DYRK1A targeting was stronger in *KMT2A*-R leukemic cells as in non-*KMT2A*-R ALL cells, 2) DYRK1A is not a common essential gene assessed via the Cancer Dependency MAP, and 3) DYRK1A has not been studied specifically in *KMT2A*-R ALL. Previous studies have suggested involvement of DYRK1A for Down syndrome (DS-ALL) associated megakaryoblastic leukemia^16^ and normal hematopoiesis^17,52^. More recently, additional studies support a critical role of DYRK1A in DS-ALL^41^. While higher DYRK1A expression levels are expected in ALL cells from patients with Down syndrome, given that DYRK1A is located on chromosome 21, it is not known how DYRK1A is transcriptionally regulated. Here we demonstrate a direct transcriptional regulation of DYRK1A via KMT2A fusions. In addition, we report that pharmacologic inhibition of Menin, an essential transcription factor involved in the regulation of *KMT2A*-R target genes^53–55^ and direct DYRK1A inhibition results in cell cycle arrest of *KMT2A*-R ALL. Given that DYRK1A is not the only target gene that is regulates by menin, we specifically focused to study the importance of DYRK1A for *KMT2A*-R ALL survival using DYRK1A inhibitors. These results demonstrate that *KMT2A*-R ALL requires DYRK1A for normal cell proliferation and inhibition of DYRK1A and BCL2 decreases progression in *KMT2A*-R B-ALL PDX models.

ALL cells are bone marrow derived B cell precursors that have lost the ability to differentiate but retain some properties of B cells. In B cells it has been shown that the PI3K and ERK signaling pathways are regulated via the B cell receptor and, depending on the signaling strength, may mediate negative selection to prevent autoimmunity. Previous results in normal B cells have suggested that DYRK1A regulates B cell differentiation^17^. Additionally, it has been demonstrated that DYRK1A regulates B cell survival and B cell autoimmunity via activation of NF-κB^52^. Our results demonstrate that DYRK1A inhibition in *KMT2A*-R ALL prevented cell proliferation while interestingly resulting in hyperphosphorylation of ERK in contrast to prior reports in other cell types^43,45^. Dual inhibition of DYRK1A and the ERK regulating kinases MEK1/2 prevented ERK hyperphosphorylation and consequent rescued cell proliferation. demonstrating that DYRK1A mediate hyperphosphorylation of ERK is specifically negatively affecting cell proliferation and survival. Given that *KMT2A*-R ALL cells do not express a (pre) B cell receptor on the surface, we hypothesize that the mechanism of negative selection is induced independent of the B cell receptor in these cells. Overall, we hypothesize that DYRK1A downregulates negative selection of *KMT2A*-R B-ALL allowing these malignant cells to rapidly proliferate. Upon DYRK1A inhibition, ERK hyperphosphorylation may stimulate a type of negative selection that makes cells vulnerable to inhibition of BH3 based cell survival mechanisms.

Another important observation is that DYRK1A and MYC negatively regulate each other in a feedback loop. First, we identified that DYRK1A deletion or pharmacologic inhibition results in increased MYC levels, which is in line with a recent publication demonstrating that viral overexpression of DYRK1A in non-*KMT2A*-R AML results in phosphorylation of MYC at Ser62 and Thr58 and consequently in MYC degradation^31^. A critical regulator of apoptosis is BIM, which is directly regulated via MYC. Interestingly, our data demonstrate that inhibition of DYRK1A results in upregulation of MYC and consequently in higher BIM levels, while the BIM negative regulator BCL2 is not affected. Strikingly, we demonstrate that combined DYRK1A and BCL2 inhibition synergistically killed *KMT2A*-R ALL cells and reduced the leukemia burden of mice injected with a *KMT2A*-R PDX samples.

Despite significant improvements in the treatment of patients with ALL, those with the *KMT2A*-R subtype experience inferior clinical outcomes. A particularly feared sub-type of ALL is infant leukemia which is, by definition, leukemia occurring in patients less than 12 months of age. Given that the frequency of *KMT2A*-R ALL is about 70% in infants with ALL, our study demonstrates that combined inhibition of DYRK1A in combination with BCL2 may be a novel treatment approach that could improve the clinical outcome of this particular patient population.

## Author Contributions

CH designed and directed the study, performed experiments, analyzed, and interpreted data, wrote the manuscript and provided research funding; VSSAA performed experiments; GW assisted with design and execution of CRISPR screen; JAC and JPL performed preclinical *in vivo* animal studies and assisted with provision of ALL cell lines and PDX model cells for in vitro studies; HG and TM provided and analyzed ChIP-Seq data; SL, AK, SS analyzed and provided RNA-Seq data and discussed results; XH and KMB provided the menin inhibitors MI-503 and VTP50469: RB and JDC provided consultation on DYRK1A biology, selection, and use of DYRK1A inhibitors and assisted with the preparation of the manuscript. TB provided EHT1610 for these collaborative studies. MC and SKT oversaw the study, assisted with experimental design, interpreted data, provided research funding, and edited the manuscript. All authors approved the final version of the manuscript.

## Acknowledgements

We thank Dr David Fruman at the University of California, Irvine for provision of TVA-1 cells used in these studies. This study was supported by the Rally Foundation for Childhood Cancer, The American Society of Hematology Restart research Grant, NIH/NCI K22 CA251649-01 (CH). S.S. was supported by a Scholar Award from the American Society of Hematology, the Andrew McDonough B+ Foundation, and the Leukemia Lymphoma Society Translational Research Program Grant 6624-21. R.S.B. was supported by the Hematopoiesis Training Program grant at the University of Pennsylvania (NIH/NIDDK T32 DK07780). S.K.T. was supported by the National Institutes of Health (NIH)/National Cancer Institute 1U01CA232486 and 1U01CA243072 awards, Department of Defense Translational Team Science Award CA180683P1, V Foundation for Cancer Research, the Rally Foundation for Childhood Cancer, and the SchylerStrong Foundation for Childhood Cancer Research. SKT is a Scholar of the Leukemia and Lymphoma Society and holds the Joshua Kahan Endowed Chair in Pediatric Leukemia Research at the Children’s Hospital of Philadelphia.

