## Supplement for "Pharmacologic Inhibition of DYRK1A Results in MYC Hyperactivation and ERK Hyperphosphorylation rendering *KMT2A*-R ALL Cells Sensitive to BCL2 Inhibition"

Supplement Figure 1

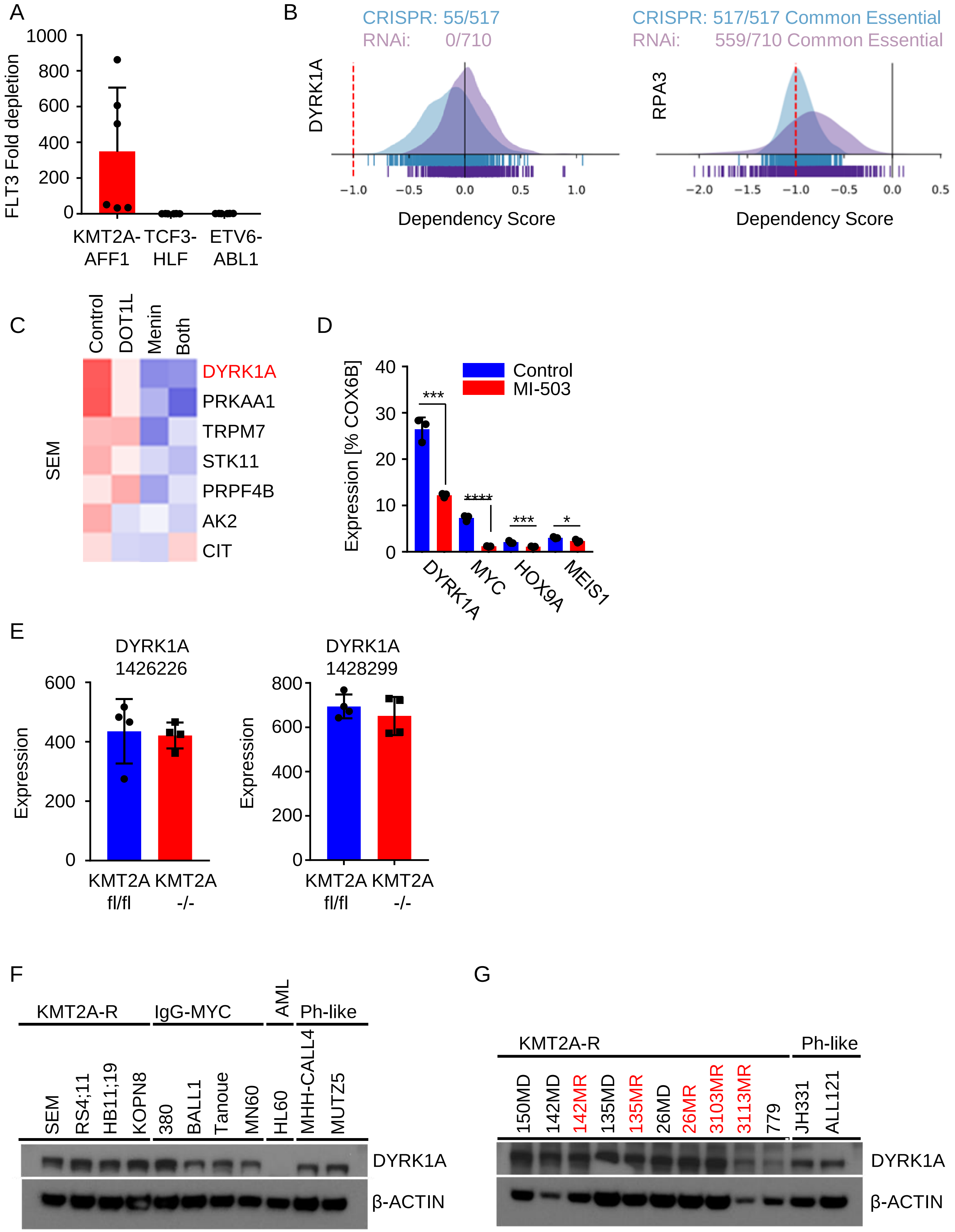

**Supplement Figure 1**

**A)** Shown is the fold depletion of FLT3 among the different ALL cell lines that were used for the kinome-wide CRISPR screen (shRNAs n=6). **B)** Dependency analysis of DYRK1A and RPA3 via the Cancer Dependency MAP website. **C)** Analysis of gene expression data from SEM cells that were treated with DMSO, 3 µM EPZ004777 (DOT1L inhibitor), 3 µM MI-2-2 (Menin inhibitor) or a combination of both for 4 days as previously published (GSE63664)[58]. Shown is a list of target genes that were identified in our kinome-wide CRISPR screen. **D)** RT-PCR was performed to determine the gene expression levels of the indicated genes in SEM cells after 5 days of treatment either with control or 3 µM MI-503 (Menin inhibitor). **E)** Gene expression analysis of DYRK1A levels after induced deletion of *KMT2A* in a conditional genetic *KMT2A* KO mouse model. Shown are the two probe sets for DYRK1A[59]. **F)** Western blot analysis of the indicated cell lines for the protein expression levels of DYRK1A and β-ACTIN. **G)** The protein levels of DYRK1A and β-ACTIN were determined via Western blot in *KMT2A*-R PDX cases at diagnosis (black) and relapse (red) two Ph-like ALL PDX cases were used as controls.

Supplement Figure 2

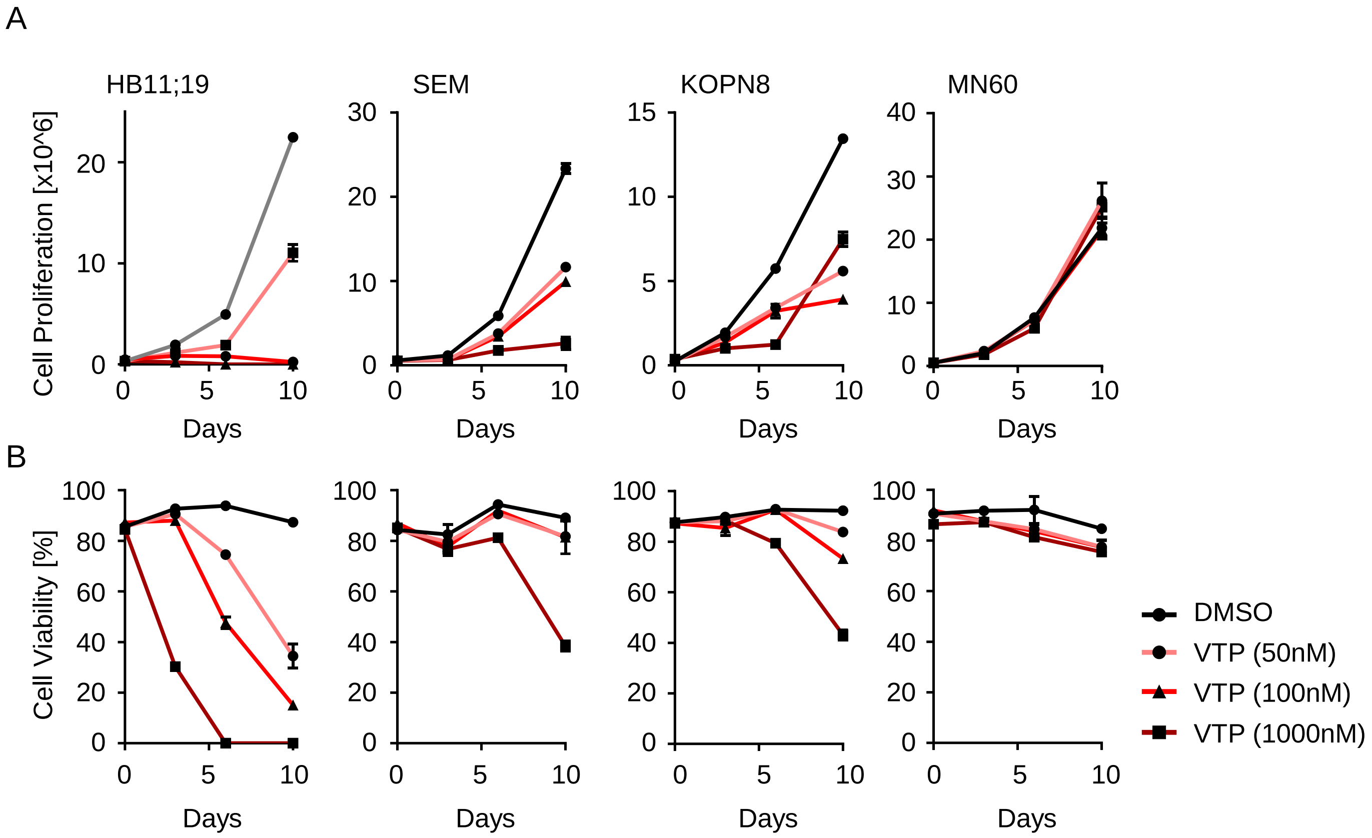


**Supplement Figure 2**
The *KMT2A*-R ALL cell lines HB11;19, SEM, KOPN8 and the non-*KMT2A*-R ALL cell line MN60 were treated with three different concentrations of the Menin inhibitor VTP50469 (50nM, 100nM, 1000nM) for the indicated time points. Cell number **(A)** and cell viability **(B)** was determined via the autonomous cell counter system (n=3).

Supplement Figure 3

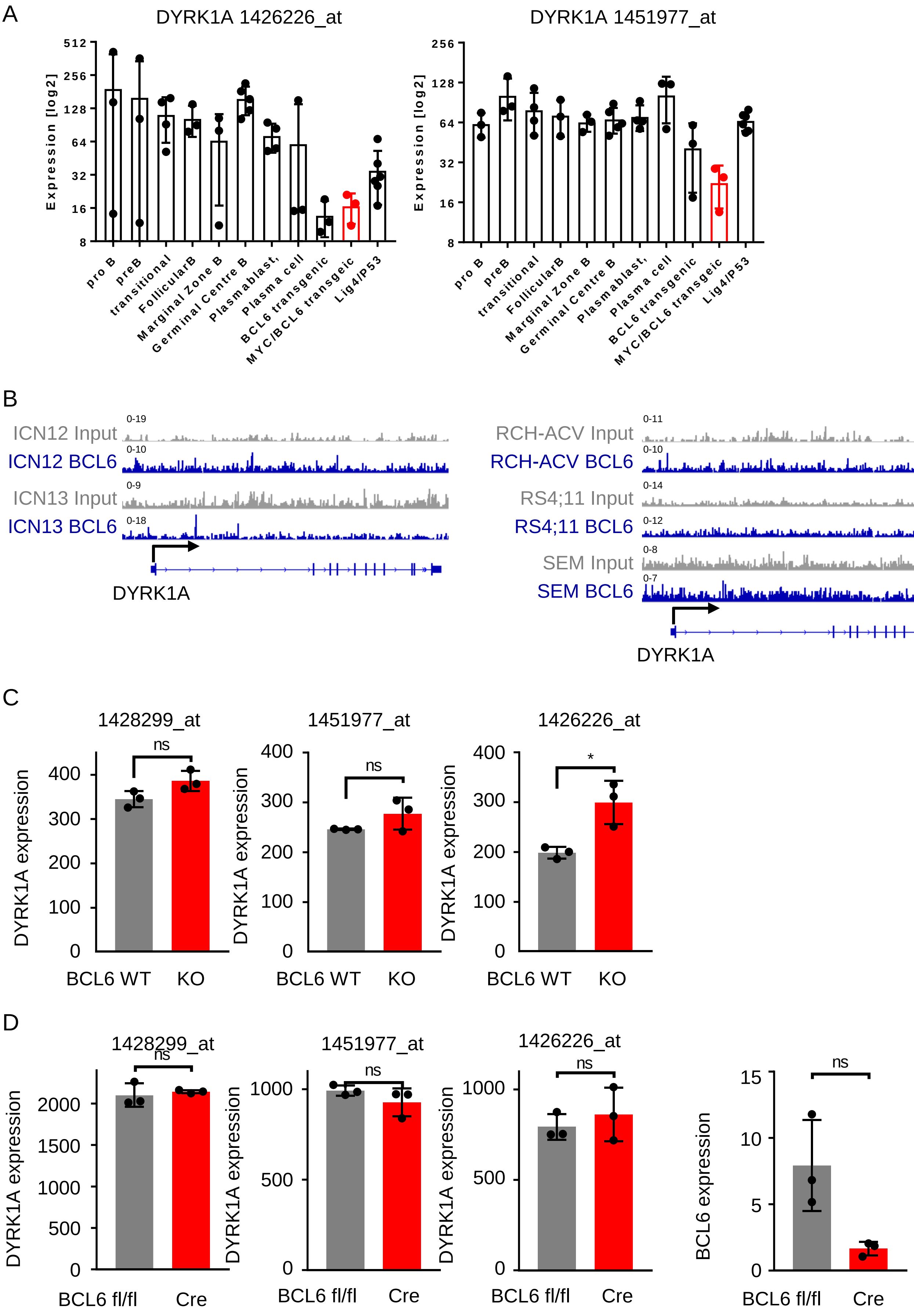

**Supplement Figure 3**

**A)** DYRK1A gene expression levels in comprehensive panel of purified developmentally defined normal murine B cells and genetically distinct murine lymphoma models (GSE26408)[34]. Shown are the probe sets 1426226_at and 1451977_at. **B)** ChIP-Seq analysis of two different ALL PDX cases (ICN12, TCF3-PBX1 and ICN13, *KMT2A*-AFF1) and three different ALL cell lines (RCH-ACV, TCF3-PBX1; RS4;11, *KMT2A*-AFF1; SEM, *KMT2A*-AFF1). **C/E)** Gene expression analyses of BCL6 WT and KO mouse ALL cells. Shown are three different probe sets for DYRK1A and one for MYC (GSE20987)[60]. **D)** Gene expression analyses of conditional BCL6$^{fl/fl}$ mouse ALL cells. Shown are three different probe sets for DYRK1A and one for BCL6. Deletion of BCL6 was induced via CRE as previously published (GSE59332)[61].

Supplement Figure 4

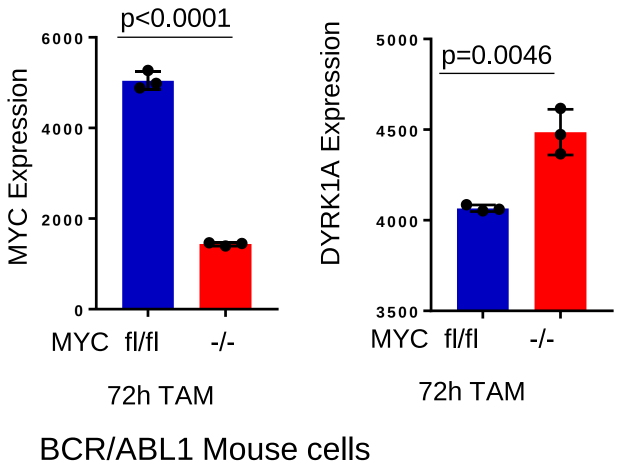


**Supplement Figure 4**

**A)** Molecular structure of Harmine. **B)** Cell proliferation/viability was determined via XTT in KMT2A-R ALL cell treated with increasing concentrations of harmine (left) and harmine hydrochloride (right). **C)** Western blot analysis of either control, harmine (5µM), or harmine hydrochloride (5µM) treated KMT2A-R ALL cell lines for 72h. Protein levels of MYC, CCND3, pERK, ERK and β-ACTIN was determined.

Supplement Figure 5

A

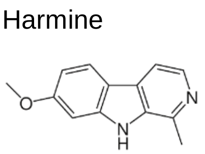


B

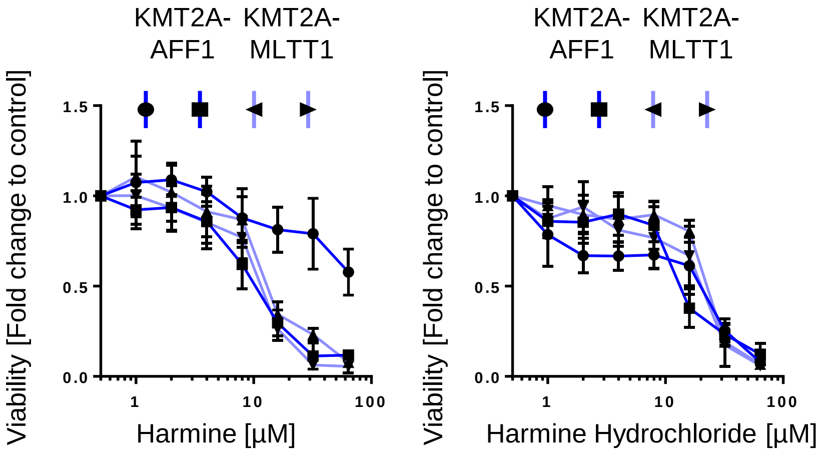


C

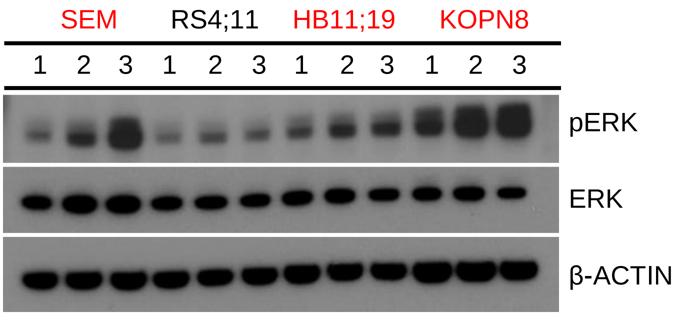


Cell lines in red have FLT3/RAS mutations
1 Control
2 Harmine 5µM
3 Harmine Hydrochloride 5µM

**Supplement Figure 5**

**A)** Gene expression analyses of conditional MYC$^{fl/fl}$ mouse ALL cells. Shown are the gene expression levels of MYC, DYRK1A and E2F1. Deletion of MYC was induced via CRE as previously published (GSE30928)[62].
